# Lead optimization against drug-resistant *Leishmania donovani* infection: Semi-synthetic derivatives of ethyl linoleate isolated from Indian edible mushroom *Meripilus giganteus*

**DOI:** 10.1101/2025.07.03.663049

**Authors:** Supriya Nath, Karan Chhetri, Aabid Hussain, Ankur Chaudhuri, Joydip Ghosh, Sondipon Chakraborty, Debarati Mukherjee, Mintu Karan, Bikramjit Raychaudhury, Krishnendu Acharya, Saikat Chakrabarti, Biswajit Gopal Roy, Chiranjib Pal

## Abstract

Visceral leishmaniasis, the most severe form, affects thousands of people annually. Current drugs in practice fail to provide an absolute cure. Globally, treatment relies on a single dose of liposomal amphotericin B, which requires a costly setup and cold chain to maintain. Natural molecules remain the vital sources of therapeutic ingredients in modern medicine. *Meripilus giganteus* is traditionally consumed by the people of the North-eastern regions of the Himalayas, India. In this study, we planned for the stepwise bioactivity-guided isolation of natural molecules from *M. giganteus* and to synthesize more potential therapeutic derivatives against the drug-resistant *Leishmania* infection. HPLC chromatogram of ethyl acetate extract obtained from *M. giganteus* revealed the presence of ethyl linoleate as the most active anti-leishmanial molecule. Several derivatives of ethyl linoleate were synthesized and evaluated against *Leishmania* promastigotes. Among them, a single epoxygenated variant of ethyl linoleate, ethyl (Z)-8-(3-(oct-2-en-1-yl) oxiran-2-yl) octanoate (EL3), showed promising anti-leishmanial activity against both drug-sensitive and drug-resistant *Leishmania donovani*. The lead derivative disrupts the biosynthesis of trypanothione from glutathione and spermidine, making the parasite vulnerable to the oxidative burst inside the host. Interestingly, the lead derivative was found to be more efficient against the drug-resistant *L. donovani*, *in vivo*. Besides this, it has almost no toxicity towards host cells. Considering the urgency for new drug development, we propose this novel semi-synthetic derivative as a potent therapeutic lead.

## Introduction

Infectious diseases cause large-scale mortalities despite remarkable discoveries and advancements in modern medical sciences worldwide. Vector-borne infectious diseases like leishmaniasis lack effective vaccines and hence pose a serious threat of major outbreaks with increasing incidence every year. Visceral leishmaniasis (VL), the deadliest form of leishmaniasis, accounts for an estimated 50000-90000 new cases annually, worldwide (1). In the year 2023, several new cases of VL were reported to the WHO from India, Ethiopia, Kenya, Somalia, South Sudan, China, Brazil, Eritrea, and Yemen, and this accounted for only 25 to 45% of actual cases that occurred across the globe (2). Certain chemotherapeutics hold the gates but lack absolute remission for many reasons. Long-term treatment regimens and irregular patient follow-ups lead to the emergence of resistant strains. Several instances of drug resistance have already been reported for sodium antimony gluconate (SAG) from the Indian state of Bihar (3). Drug resistance in *Leishmania* results from a high rate of drug efflux attributed to the over-expression of membrane-associated proteins like ABC transporter-MRPA or due to the scavenging and pro-parasitic activities of the enzymes of polyamine biosynthetic pathways (4). Liposomal Amphotericin B, the present first-line chemotherapy against VL, is not cost-effective (5). Considering all these drawbacks, we propose naturally occurring purified products and their derivatives as potential candidates for future drug discovery. Previously, we have shown that a triterpenoid, astrakurkurone, isolated from *Astraeus hygrometricus*, could inhibit the proliferation of *L. donovani* and protect the host from experimental VL by inducing immunity (6, 7). A semi-purified carbohydrate fraction from the same mushroom showed an anti-parasitic response by inducing pro-inflammatory cytokines (TNF-α, IL-12, iNOS2) in hosts (8). So, mushrooms can be a good source of potential anti-inflammatory molecules; extracts were previously cited for their anti-viral, anti-bacterial efficacy (9). In the present study, we aimed to isolate active principles from the wild mushroom *M. giganteus* and synthesise chemical derivatives with the aim of drug development against both the drug-sensitive (10, 11) and the drug-resistant *L. donovani* infection (12,13). Based on our preliminary observation, a semi-purified ethyl acetate fraction of *M. giganteus* was found to show a high inhibitory effect against *L. donovani* (Fig. S1B), it was further purified through HPLC, resulting in six distinct peaks on the HPLC chromatogram (Fig. S1D). Peak 5, characterized as ethyl linoleate, was found most efficient against the promastigotes of drug-sensitive and drug-resistant *L. donovani* (Fig. S1E). Targeting for better anti-leishmanial efficacy, chemical derivatization of ethyl linoleate (Peak 5) was performed by controlled oxygenation that yielded five derivatives, (EL2-EL6) (Fig. 1). Interestingly, monoepoxides of ethyl linoleate (EL3) were found to inhibit the proliferation of both promastigote and amastigote morphs of drug-sensitive as well as drug-resistant *L. donovani*, *in vitro*, most significantly (Fig. 2, 3, Table S3, S4, S7A, B).

**Figure 1:**
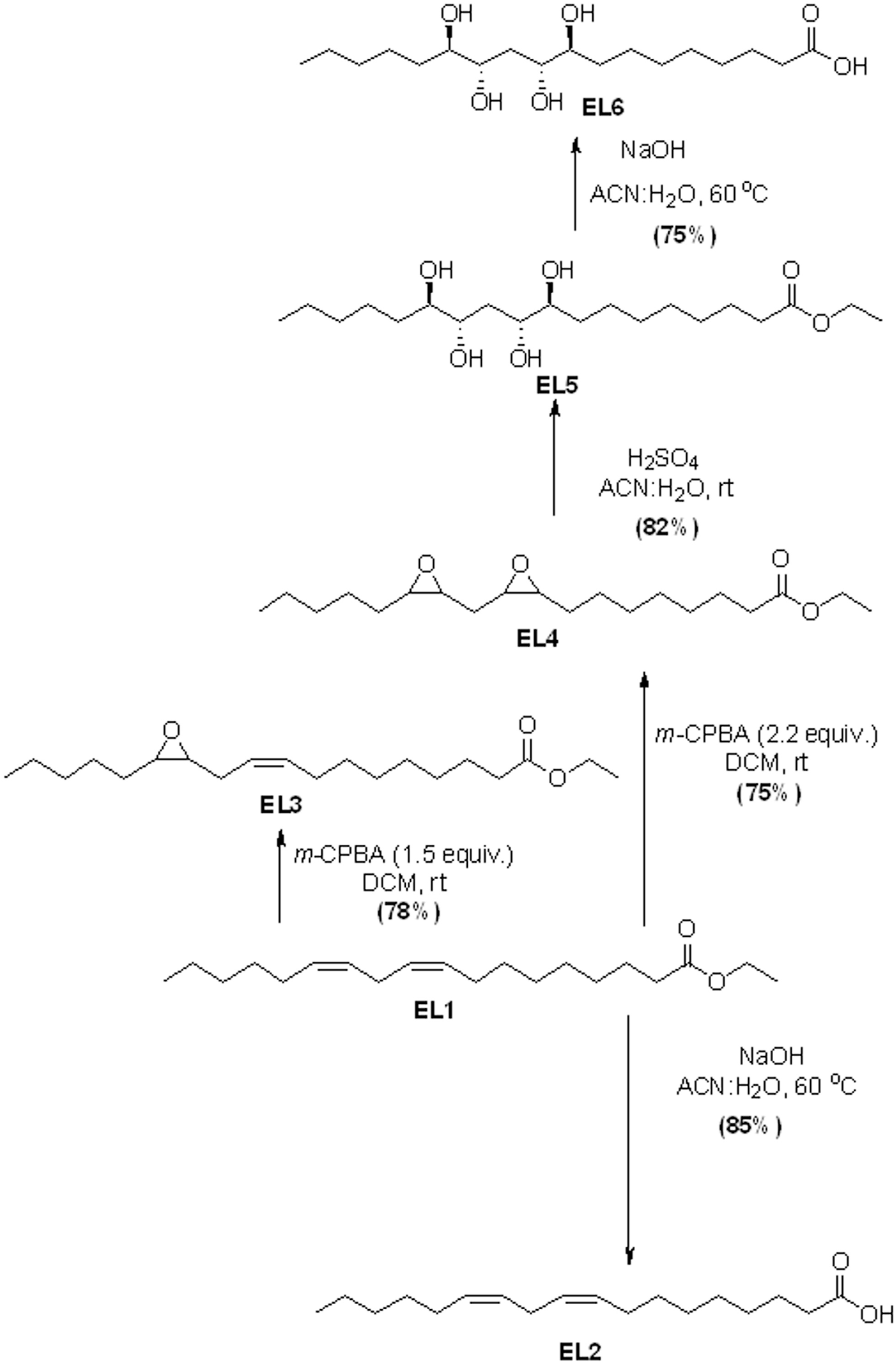
Synthesis scheme of the derivatives (EL2-EL6) from Peak 5/ EL1.

**Figure 2:**
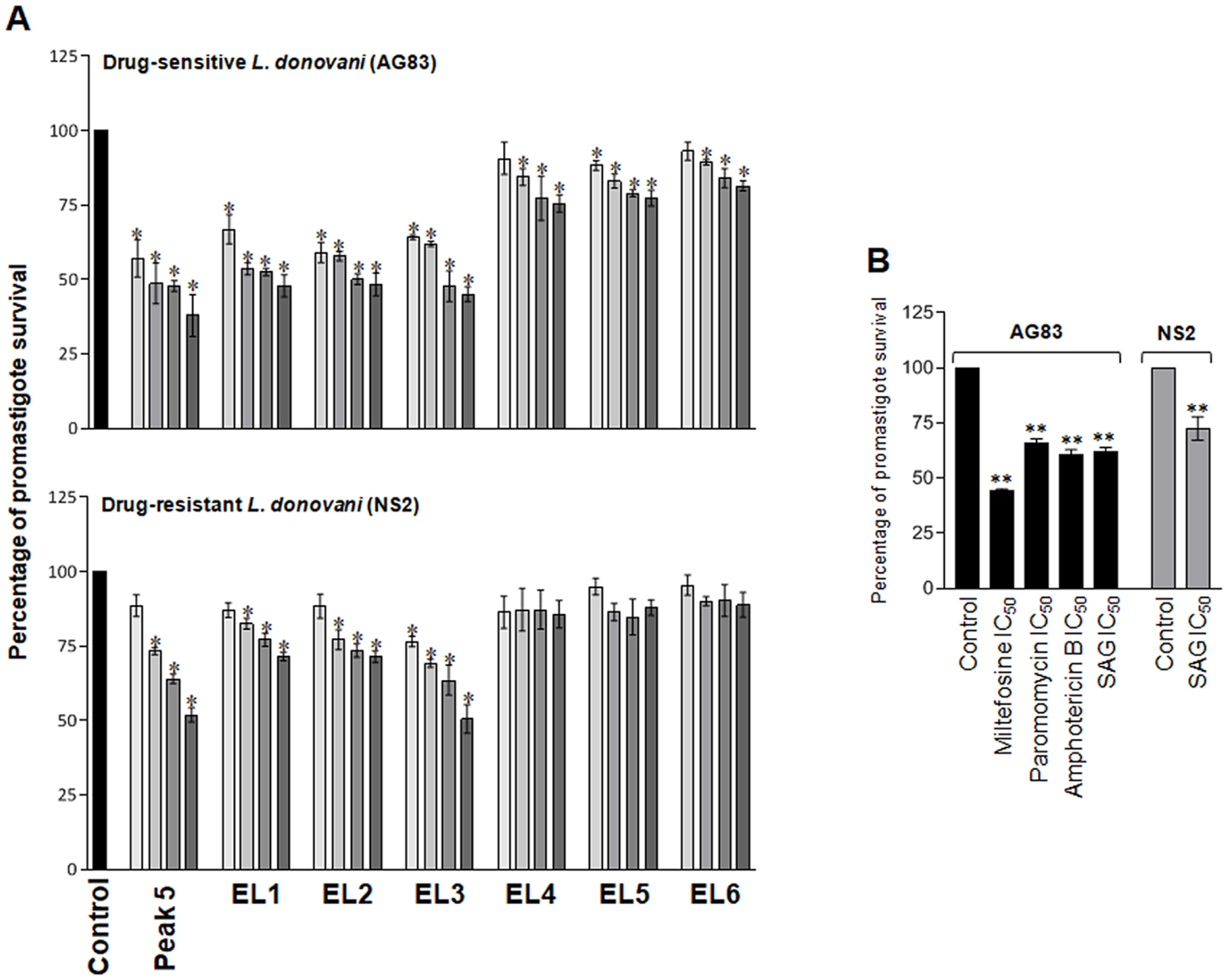
Anti-promastigote activity of ethyl linoleate and synthetic derivatives, *in vitro*. (A) Original Peak 5 compound and EL3 reduced the proliferation dose dependently as observed by MTT assay. Results are shown as mean ± SEM of three different experiments performed in triplicate; *p<0.05 vs. DMSO-treated control. (B) The IC50 doses of reference drugs were tested on both AG83 and NS2 parasites, **p<0.001 vs. control.

The background study was intended for us to focus on some questions to identify EL3 as a promising anti-leishmanial lead. The first question was the route of administration and the dose selection. The second question was the mechanism of action; does EL3 have any specific target on the intracellular parasites in the host? Does it induce the pro-inflammatory responses of the hosts, leading to parasite clearance? The third question was regarding the toxicity towards hosts, if any. Finally, the bioavailability of EL3 in the host system was investigated, as the longer persistence of drug molecules inside the host system promotes resistance.

## Results

### Isolation of active molecules from *Meripilus giganteus*

Among the different solvent-extracted fractions, the significant *in vitro* bioactivity of the ethyl acetate-extracted fraction against the drug-resistant strain of *Leishmania* (Fig. S1C) prompted us to bioactivity-guided isolation of pure molecules from this fraction. Based on retention times in the optimized mobile phase, six distinct peaks were isolated from the bioactive fraction of the ethyl acetate extract, as depicted in the chromatogram (Fig. S1D), through multiple injections and run sequences using a semi-preparative column. Bioactivity assays against *L. donovani* promastigotes indicated that fraction 5 exhibited the highest activity. After confirming their bioactivity (Fig. S1E), we proceeded with the structural determination of the active compound (s). Simple ¹H and ¹³C NMR spectral analysis confirmed that, as anticipated, the active fractions consisted of pure single compounds rather than mixtures.

The structure of active compounds obtained from peak 5 was determined by thorough analysis of infrared, mass spectrometric (HRMS), and nuclear magnetic resonance (1D and 2D NMR) spectroscopic data. A quartet peak at δ 4.15 ppm for two protons (2H) and a triplet peak for three protons (3H) at δ 0.9 ppm in ^1^H NMR spectrum of the pure compound obtained from peak 5 suggested the presence of an ethyl ester. The carbon peak at δ 174.5 ppm in ^13^C NMR spectra supported the presence of ester carbonyl carbon (Fig. S2). The presence of a strong, sharp carbonyl peak at 1735 cm^-1^ in the IR spectrum further strongly supports the presence of the ester group. The peak at δ 5.3 ppm in ^1^H NMR for four protons (4H) suggested the possibility of the presence of multiple similar double bonds with similar chemical environments, which are probably in coupling with multiple aliphatic protons. The appearance of multiple protons in ^1^H NMR spectrum and multiple methylene (-CH_2_) peaks at DEPT 90 spectrum indicated the presence of multiple methylene groups in the molecule (14). The correlations among protons in the molecule in correlation spectroscopy (COSY) and C-H correlation data obtained from the HMQC spectrum of the compound suggest that the compound was a long-chain fatty acid ester with two isolated double bonds (15,16). Space integration in NOESY among allylic protons suggests that these double bonds are in cis orientation (17). All the above observations, along with the electrospray ionized mass spectrometric (ESI) molecular ion peak at m/z 308.2725, corresponding to the molecular formula C_20_H_36_O_2_, led us to determine the structure of the isolated compound in peak 5 as ethyl linoleate EL1, a known di-unsaturated long-chain fatty acid ester (Fig. S2). The determined structure of the identified compound was further confirmed by comparing it with the reported proton and carbon NMR spectra of ethyl linoleate.

### Synthesis of a library of polar derivatives of EL1 with increased aqueous solubility

As EL1 (Fig. S2) is a di-unsaturated ester, we could make it comparatively polar by hydrolyzing it to linoleic acid EL2 (Fig. S3), through base-catalyzed hydrolysis using sodium hydroxide solution, without affecting the olefins present in the molecule (18). The successful hydrolysis of EL1 is evident from the disappearance of the characteristic ethoxy (-OEt) quartet peak at δ 4.12 ppm for two protons (2H) and the triplet peak at δ 0.90 ppm for three protons (3H), in ^1^H NMR spectra of product EL2. Considering dihydroxylation of the existing double bonds of EL1 could be an effective approach to generate more polar derivatives with enhanced aqueous solubility, we planned to carry out dihydroxylation of both the existing double bonds. However, when we tried to carry out cis-hydroxylation of both the double bonds of EL1 using osmium tetroxide and *N*-methyl morpholine oxide, it failed to react to generate the desired tetrahydroxylated product (19). The failure of the direct di-hydroxylation method using osmium tetroxide on EL1 forced us to look for an alternative approach. Our endeavor to epoxidize the olefins of EL1 using 1.5 and 2.5 equiv. of meta-chloroperbenzoic acid (*m*CPBA) in dichloromethane (DCM) at room temperature, successfully resulted in the formation of mono-epoxy EL3 (Fig. S4) and di-epoxy derivative EL4 (Fig. S5) (20). Confirmation of the formation of monoepoxide EL3 was evident from the reduction in the number of olefinic protons at δ 5.38 ppm from four (in the case of EL1) to two protons and an upfield shift of two protons to δ 2.91 ppm due to the characteristic anisotropic shielding of epoxide. Similarly, the confirmation of the formation of EL4 was asserted from the disappearance of all four olefinic proton peaks and the upfield appearance of these protons at δ 3.24–3.04 and 2.98 ppm as two multiplets due to the epoxide anisotropy.

The diepoxide EL4 was then subjected to acid-catalyzed hydrolysis using dilute sulfuric acid (4%) in an acetonitrile/water (1:1) mixture, yielding the tetrahydroxylated long-chain ethyl ester EL5 (Fig. S6). The formation of EL5 was supported by the expected downfield shifts of the epoxide-associated protons from δ 3.24–3.04 and 2.98 ppm in EL4 to δ 3.30–4.3 ppm in the ¹H NMR spectrum of EL5. To further enhance polarity, we hydrolyzed the ester group of EL5 using aqueous NaOH, converting it into the corresponding tetrahydroxy acid EL6 (Fig. S7). The successful ester hydrolysis was confirmed by the selective disappearance of the ethoxy peak in EL5, indicating the formation of EL6. ^1^H and ^13^C NMR spectra, structures, and synthesis of all derivatives have been provided in the supplementary file (Fig. S3-S7).

### Activity-guided selection of the bioactive synthetic derivative against *L. donovani* promastigotes, in vitro

EL1 (IC_50_ for AG83 promastigotes, 80.3 ± 2.6 µg/ml, p<0.05 vs. DMSO treated control; IC_50_ for NS2 promastigotes, 176.1 ± 4.4 µg/ml, p<0.05), EL2 (IC_50_ for AG83 promastigotes 79.4 ± 2.9 µg/ml, p<0.05; IC_50_ for NS2 promastigotes 161.9 ± 5.9 µg/ml, p<0.05), and EL3 (IC_50_ for AG83 promastigotes, 77.6 ± 3.2 µg/ml, p<0.05; IC_50_ for NS2 promastigotes, 99.8 ± 5.7 µg/ml, p<0.05), vs. DMSO treated control, have been identified among the five derivatives (EL2-EL6) of Peak 5/ EL1 as these synthetic derivatives exhibited potential anti-promastigote activity (Fig. 2A, Table S3-S5). Interestingly, the rate of inhibition by EL3 was found higher against the drug-resistant NS2 (49.5±2.8% of inhibition by highest dose, p<0.05 vs. control) when compared with the Peak 5 (48.1±1.4%, (p<0.05 vs. control for NS2), originally obtained from *M. giganteus, or* EL1 (28.5±0.8%, p< 0.05 vs. control for NS2), a synthetic analogue of Peak 5 (Fig. 2A).

### EL3 showed significant anti-amastigote activity with the least toxicity, *in vitro*

EL1, EL2, and EL3 were chosen for further experiments for anti-leishmanial screening against the amastigotes, the pathogenic morphs in mammalian hosts. The dose kinetics experiment against the drug-sensitive amastigotes confirmed that EL3 (IC_50_: 62.9±2.5 µg/ml, p<0.001 vs. control) was the most efficient in inhibiting the amastigotes in infected murine peritoneal macrophages in comparison to Peak 5 (IC_50_: 87.7±1.4 µg/ml, p<0.001 vs. control), EL1 (IC_50_: 90.1±3.3 µg/ml, p<0.001 vs. control), and EL2 (IC_50_: 132.5±1.4 µg/ml, p<0.001 vs. control). EL3 was also shown to have the most significant anti-amastigote activity against the drug-resistant parasites (IC_50_: 43.8±0.6 µg/ml, p<0.001 vs. control) in comparison to Peak 5 (IC_50_: 87.7±1.3 µg/ml, p<0.001 vs. control), EL1 (IC_50_: 94.4±1.9 µg/ml, p<0.001 vs. control), and EL2 (IC_50_ 65.7±5.8 µg/ml, p<0.001) (Fig. 3A and Table S7A, B). As EL3 has been identified as the most promising synthetic derivative*, in vitro*, we have progressed with further experiments, *in vivo*, leading to the development of a successful anti-leishmanial lead.

**Figure 3:**
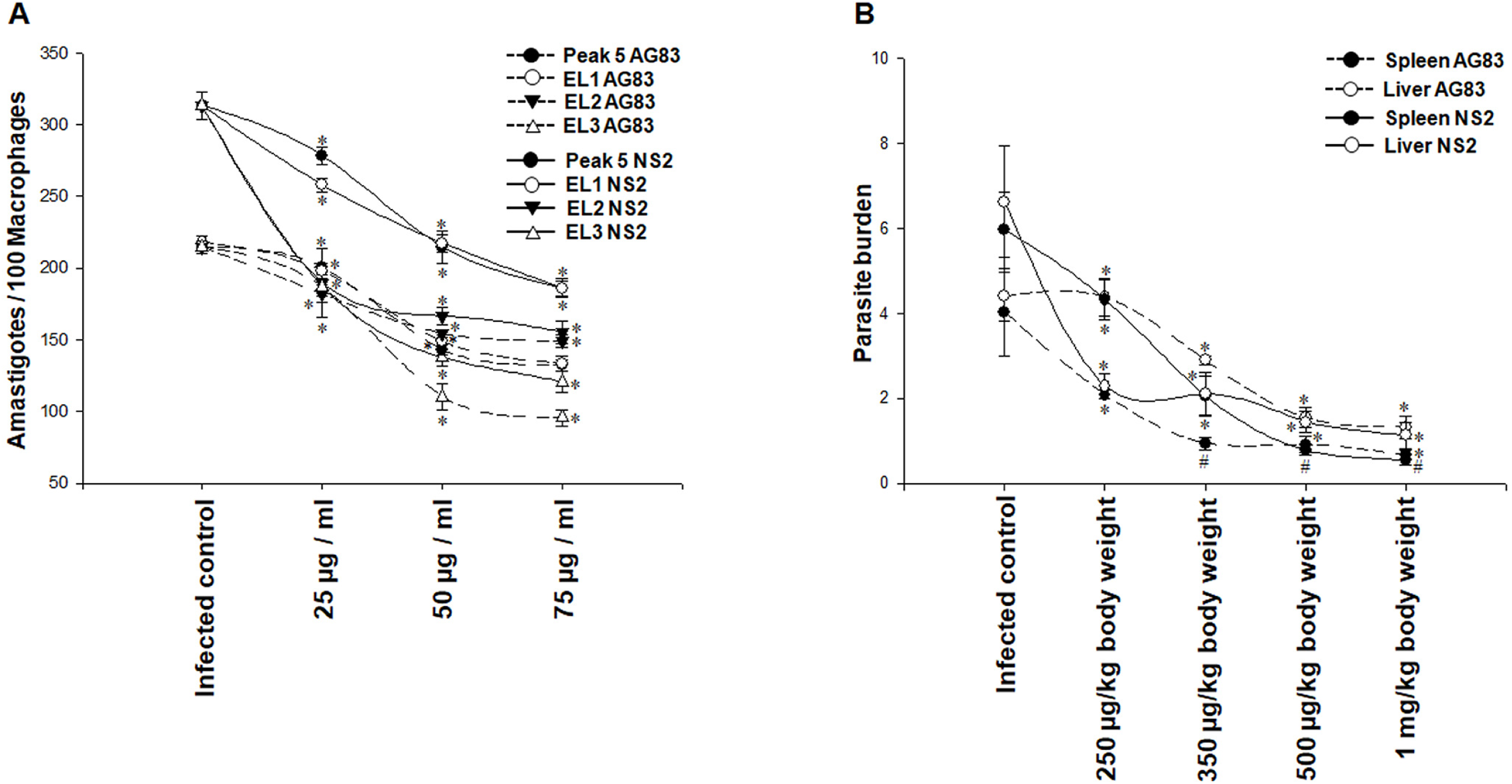
Anti-amastigote activity of ethyl linoleate and synthetic derivatives, *in vitro* and *in vivo*.(A) Anti-amastigote activity of EL1, EL2, and EL3 against intracellular amastigotes in infected macrophages, *in vitro*. (B) The anti-amastigote effect of EL3, *in vivo*. Data represented as the mean ± SEM of three different experiments performed against DMSO-treated control, and significance was calculated by ANOVA using GraphPad Prism (Version 8), *p<0.001, #p<0.05.

### EL3 reduced the drug-resistant *L. donovani* infection in visceral organs more efficiently

The epoxygenated derivative EL3 inhibited the parasite proliferation of both drug-sensitive and drug-resistant *L. donovani* infection dose-dependently, *in vivo*. The highest doses of 500 µg/kg body weight and 1mg/kg body weight EL3 were found to be more effective against the drug-resistant *L. donovani* infection in comparison to infected control and, interestingly, drug-sensitive infection (Fig. 3B and Table S9). The dose of 500 µg/kg body weight inhibited drug-resistant *L. donovani* infection by 86.7 ± 0.1% (p<0.001 vs. DMSO-treated control) and by 77.3 ± 0.3% (p<0.001 vs. DMSO-treated control) in the spleen and liver, respectively. The higher dose of 1mg/kg body weight was found to be more efficient in inhibiting the parasite proliferation by 90 ± 0.1% (p<0.001 vs. DMSO-treated control) and by 83.3 ± 0.3% (p<0.001 vs. DMSO-treated control) in the spleen and liver, respectively (Figure 3B and Table S9). This observation opens the possibility of target-specific activity of EL3 on drug-resistant parasites.

### EL3 effectively downregulates the polyamine biosynthesis pathway of parasites with a higher efficacy against drug-resistant infection

There was a significant decrease in the expression of *Leishmania* survival enzymes, trypanothione reductase (TR), glutathione synthetase (GS), and γ-glutamyl cysteine synthetase (γ-GCS) transcripts for both drug-sensitive and drug-resistant amastigotes in infected spleens with respect to normalized *L. donovani* 18s rDNA as the housekeeping gene following treatment. More interestingly, EL3 showed a consistent pattern of higher efficacy in inhibiting the expression of γ–GCS, TR, and GS mRNA of drug-resistant amastigotes when the infected animals were treated *in vivo*. The fold change of trypanothione reductase at the transcript level was found to be decreased by 50 ± 0.003-fold (p<0.001) in the case of drug-resistant amastigotes, higher than the case of drug-sensitive amastigotes (10 ± 0.06-fold, p<0.001) (Figure 4A and Table S10). In contrast, EL3 could not dampen the expression of host-specific γ-GCS, GS mRNA expressed in the infected spleen (Figure 4B and Table S11).

**Figure 4:**
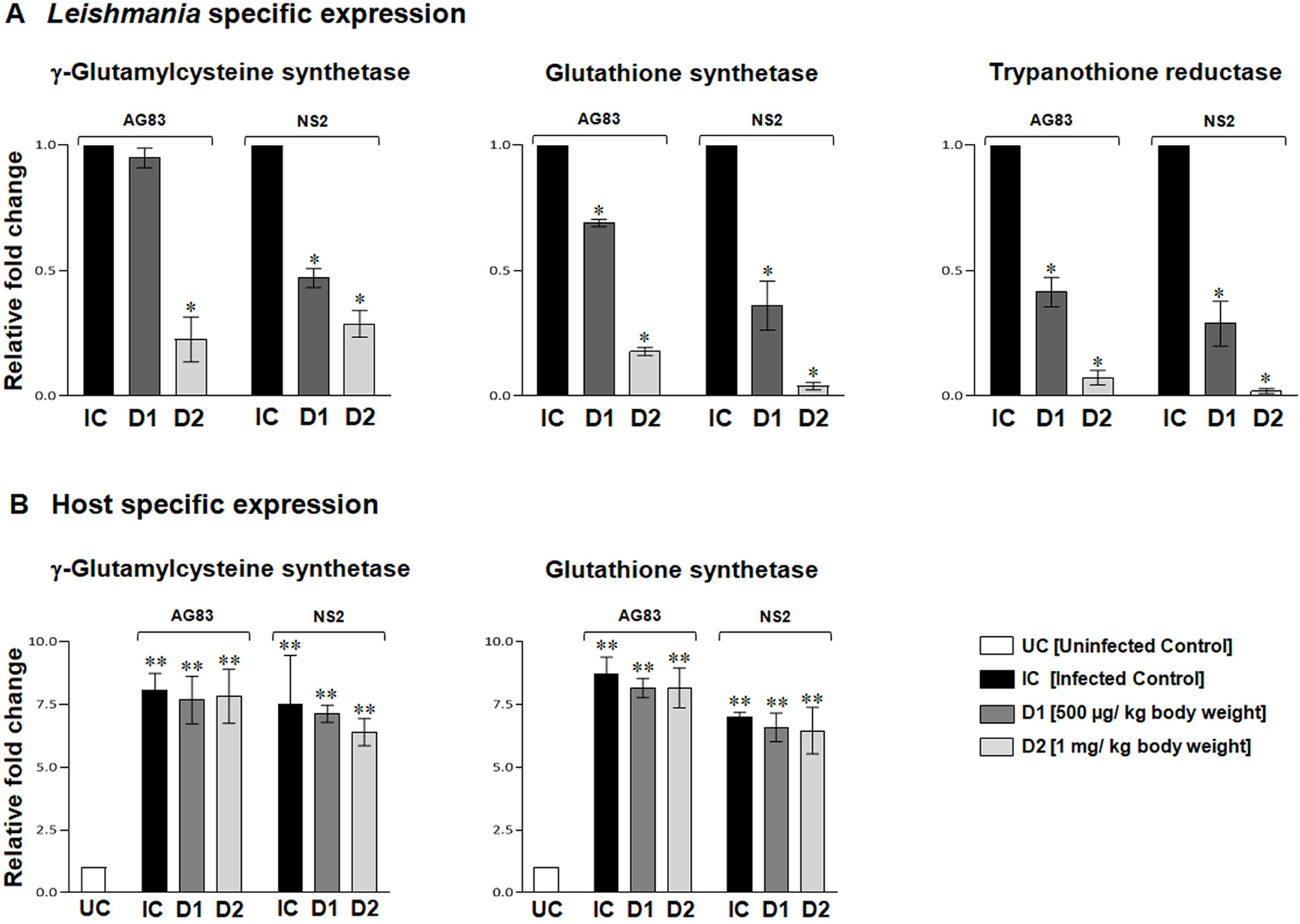
Relative mRNA expression of the polyamine biosynthesis pathway key enzymes of the parasite and the murine host. (A) The relative γ-GCS, GS, and TR (*Leishmania* specific) mRNA expressions in the spleens of experimental animals, *p<0.05. (B) The relative mRNA expressions of γ-GCS and GS (host-specific) in the spleens of experimental animals, **p<0.001. Data represented as mean± SEM of two experiments, with 5 mice in each group.

### Three-dimensional structure prediction of 3 potential targets, γ-glutamyl cysteine synthetase (γ-GCS), glutathione synthetase (GS), and trypanothione reductase (TR)

The three-dimensional structures of γ-GCS, GS, and TR from *L. donovani* were successfully generated using a combination of AlphaFold2 Colab and MODELLER. Validation with PROCHECK confirmed that the modeled structures are within acceptable stereochemical parameters, ensuring that these structural models can be reliably used for further in-depth analyses. Post-validation, the AlphaFold2-generated γ-GCS model and the MODELLER-generated models for glutathione synthetase and trypanothione reductase were selected for further studies. The AlphaFold2 model of γ-GCS showed a high confidence level, indicated by a pLDDT score of 89.9. This score indicates reliable prediction accuracy and structural confidence. In contrast, the MODELLER-generated structures for glutathione synthetase and trypanothione reductase showed significant sequence homology with the known crystal structures 2WYO (*Trypanosoma brucei* glutathione synthetase) and 2JK6 (trypanothione reductase from *L. infantum*), sharing 46.7% and 98.37% sequence identity, respectively. These sequence identities suggest a better alignment of the predicted modeled structures with experimentally validated templates, particularly for trypanothione reductase, which closely resembles its reference structure. The sequence identity and validation results underline the structural integrity of these models, which were leveraged for ligand binding studies, molecular dynamics simulations, and other functional analyses.

### Binding affinity of EL3 with γ-GCS, GS, and TR

To evaluate the effectiveness and inhibition potential of the EL3 against γ-glutamyl cysteine synthetase (γ-GCS), glutathione synthetase (GS), and trypanothione reductase (TR), molecular docking studies were conducted using LeDock. From the 100 docking conformations or poses generated for each protein-EL3 complex, the lowest binding energy pose was selected for intermolecular interaction analysis. The results demonstrated that the EL3 exhibited binding energies of −3.15 kcal/mol, −3.52 kcal/mol, and −4.52 kcal/mol with γ-GCS, GS, and TR, respectively. The negative binding energy values indicate that EL3 can effectively bind within the receptor cavities of γ-GCS, GS, and TR. For γ-GCS, the EL3 interacted through hydrophobic contacts with residues Asp470 and Gln473, formed a hydrogen bond with Arg494, and established a salt bridge with Lys483, highlighting its affinity for the binding site (Fig. 5A). In GS, the EL3 binding was stabilized primarily by hydrophobic interactions with Ile172, Leu491, and Val587 (Fig. 5B). In TR, hydrophobic interactions were noted with Val34, Val46, Thr51, Thr334, and Ala337, facilitating ligand binding within the binding site (Fig. 5C).

**Figure 5:**
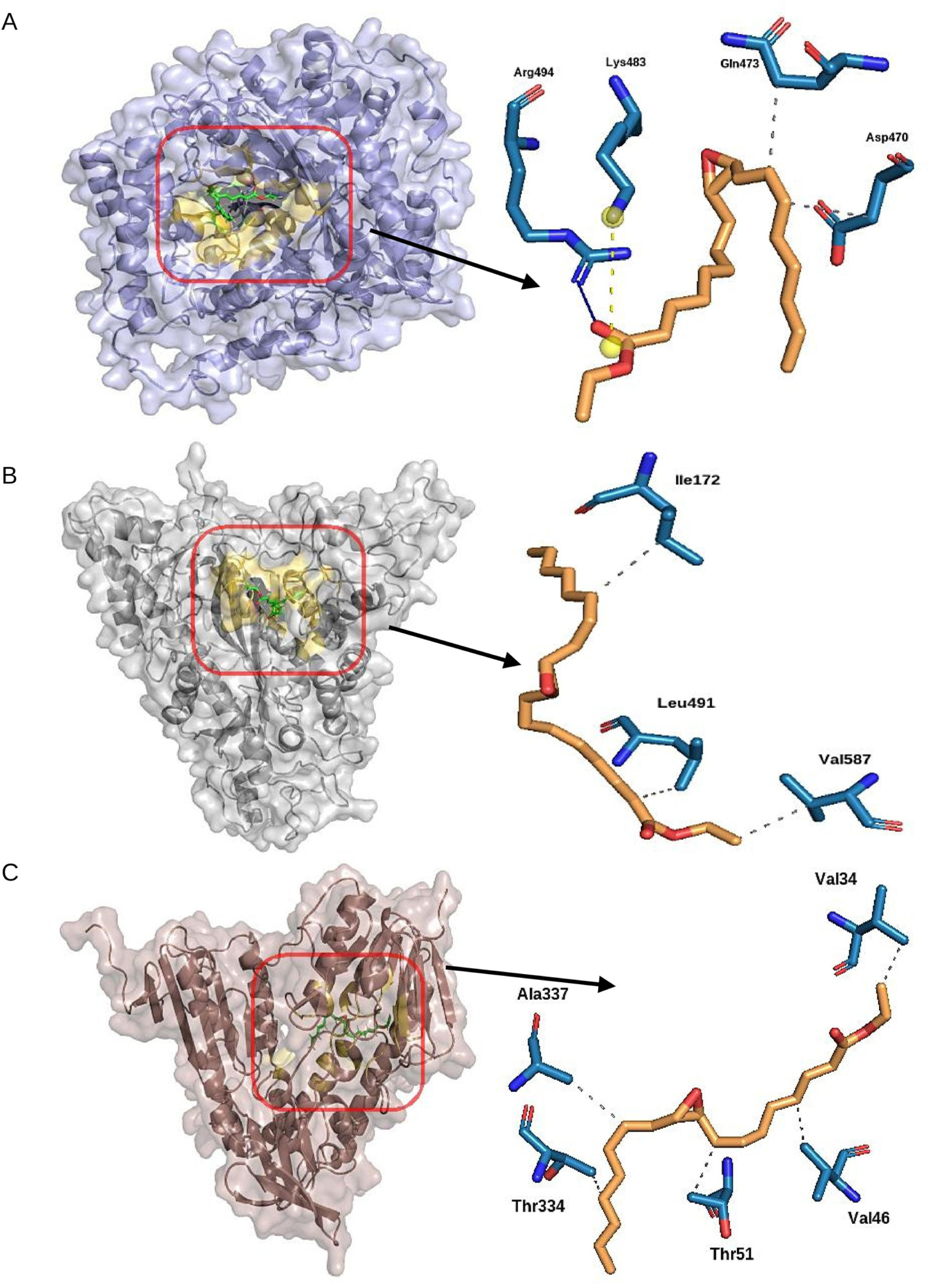
Molecular interactions of EL-3 with three target proteinsγ-glutamyl cysteine synthetase, glutathione synthetase, and trypanothione reductase. Each protein is depicted in a combination of surface and ribbon structures, while EL3 is shown in stick representation. A red box highlights a close-up view of the binding site, which is visualized as a yellow area within a 5 Å radius from EL3. (A) γ-GCS-EL3 complex. Blue, yellow, and grey lines (solid and dotted) represent hydrogen bonds, salt bridges, and hydrophobic interactions. (B) GS - EL3 complex. Grey dotted lines represent hydrophobic interactions. (C) TR - EL3 complex. Grey dotted lines represent hydrophobic interactions.

### MD simulations of three protein-EL3 complexes

The molecular dynamics (MD) simulation results for γ-GCS, GS, and TR over a 100 ns simulation are illustrated in graphical format. The energy value indicates that all proteins were stabilized quickly, with GS achieving the lowest energy, suggesting a more stable or lower-energy conformation (Fig. 6A). The root-mean-square deviation (RMSD) plot shows the stability of each protein-EL3 complex, where γ-GCS and TR maintain relatively stable RMSD values of 0.38 nm and 0.91 nm, respectively, indicating minimal structural deviations. However, GS exhibits higher RMSD deviations, reaching an average of 1.91 nm, suggesting greater structural instability in the ligand binding region throughout the simulation time (Fig. 6B). The root-mean-square fluctuation (RMSF) aligns with these observations; GS shows more prominent fluctuations across amino acid residues with an average RMSF of 0.22 nm, compared to the lower values observed for γ-GCS (0.09 nm) and TR (0.12 nm) (Fig. 6C).

**Figure 6:**
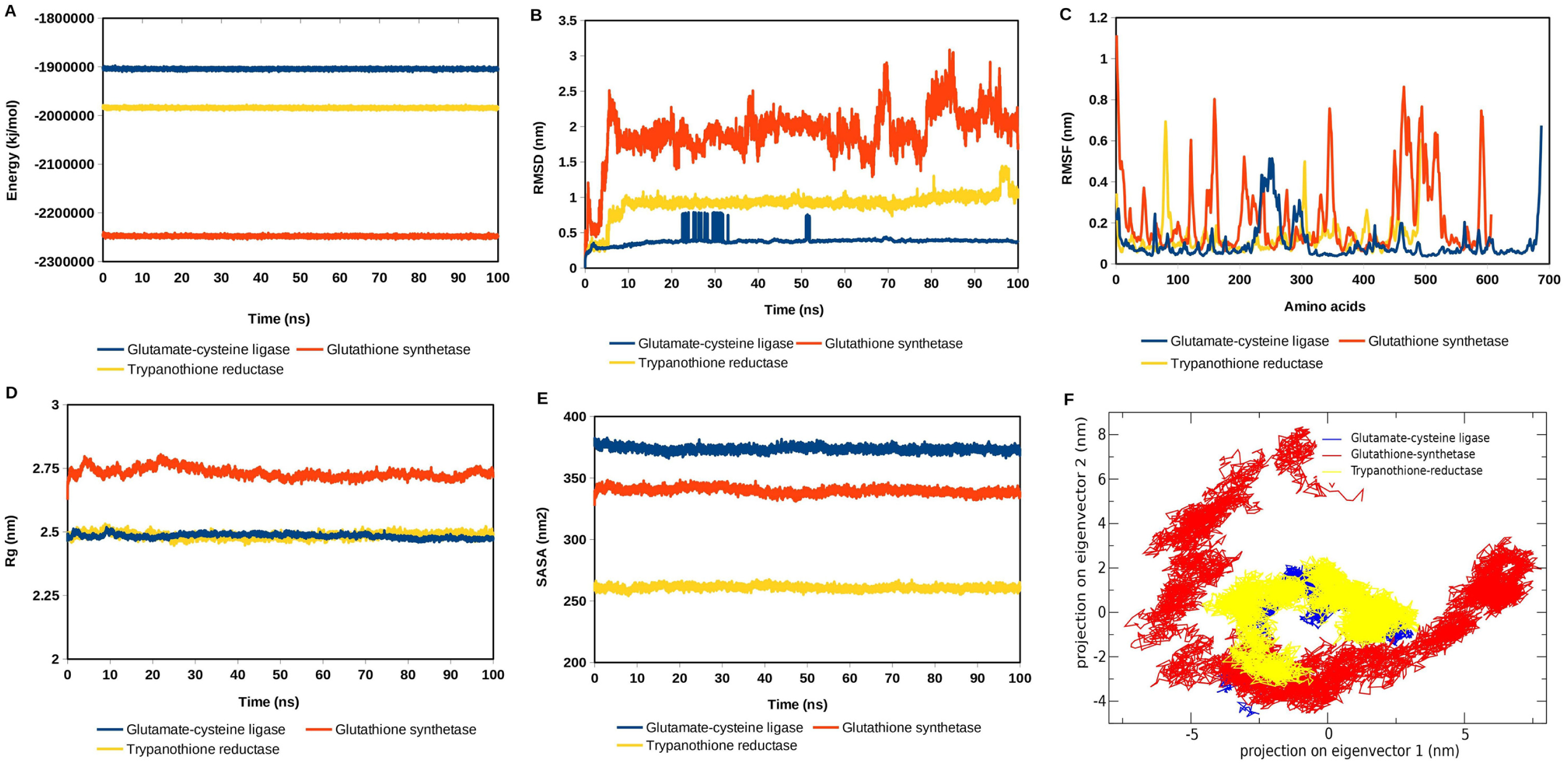
MD simulation trajectory parameters of three protein-ligand complexes. (A) Energy; (B) RMSD; (C) RMSF; (D) Rg; (E) SASA; (F) Trace of covariance matrix (PCA). RMSD: Root mean square deviation; RMSF: Root mean square fluctuation; Rg: Radius of gyration; SASA: Solvent-accessible surface area. γ-GCS, GS, and TR were represented as blue, red, and yellow lines, respectively.

The radius of gyration (Rg) values reveals the compactness of the proteins, with γ-GCS and TR both averaging 2.48 nm, signifying stable, compact structures, while GS has a slightly higher Rg value of 2.73 nm, indicating a less compact structure (Fig. 6D). The solvent-accessible surface area (SASA) results further indicate distinct protein-EL3 exposure to the solvent environment, with glutathione synthetase having the highest surface area value (339.63 nm²), suggesting increased solvent interaction, whereas γ-GCS and TR show lower SASA values of 373.64 nm² and 260.89 nm², respectively (Fig. 6E).

Essential dynamics (ED) refers to the application of principal component analysis (PCA) to a protein trajectory, allowing the extraction of essential motions from the movement of the protein molecule. This approach was employed to explore the conformational space of the three target proteins. In the principal component analysis, the trace of covariance values reflects the overall motion within the complexes. GS shows the largest trace (51.97 nm²), implying extensive conformational changes, whereas γ-GCS and TR have much smaller traces (12.06 and 11.11 nm², respectively), consistent with more restricted movements (Fig. 6F). These results collectively demonstrate that GS exhibits the highest flexibility and structural variation, while γ-GCS and TR display more stable conformations under MD conditions. This variation in structural dynamics could be relevant to the binding affinities and functional roles of these proteins.

### Comparative binding free energy study by MM-PBSA

The binding free energy calculations for the EL3 ligand with γ-GCS, GS, and TR yielded values of −12.31±7.92 kcal/mol, −15.41±3.63 kcal/mol, and −29.56±9.28 kcal/mol, respectively, indicating varied binding affinities across the three target proteins. Among the three complexes, TR exhibited the most favorable binding energy, suggesting a strong and stable interaction with EL3, while γ-GCS had the least favorable binding energy. The lower binding energy for TR may be attributed to more optimal binding interactions, likely resulting in a more stable protein-ligand complex relative to the others. The intermediate binding free energy observed for GS reflects a moderate affinity for EL3, which is stronger than that of γ-GCS but still less favorable than TR.

### EL3 did not display any hepato- or nephrotoxicity in the hosts

Evaluation of toxicity, post-treatment *in vivo,* was assessed through the estimation of serum creatinine and aspartate transaminase (AST), alanine transaminase (ALT), and alkaline phosphatase (AP) levels. Serum creatinine levels spiked in the infected, untreated control animals (1.3 ± 0.07mg/dL, p<0.001 for AG83 IM and 1.67 ± 0.21mg/dL, p<0.001 for NS2 IM) as compared to the uninfected group (0.83 ± 0.02mg/dL). However, creatinine level was reduced greatly and found to align with the normal range-0.8 ± 0.11 mg/dL for AG83 and 0.79 ± 0.07mg/dL for NS2-infected mice by the 1mg/kg body weight dose. Similar patterns were also observed in AST, AP, and ALP levels in a dose-dependent fashion (Fig. 7 and Table S12).

**Figure 7:**
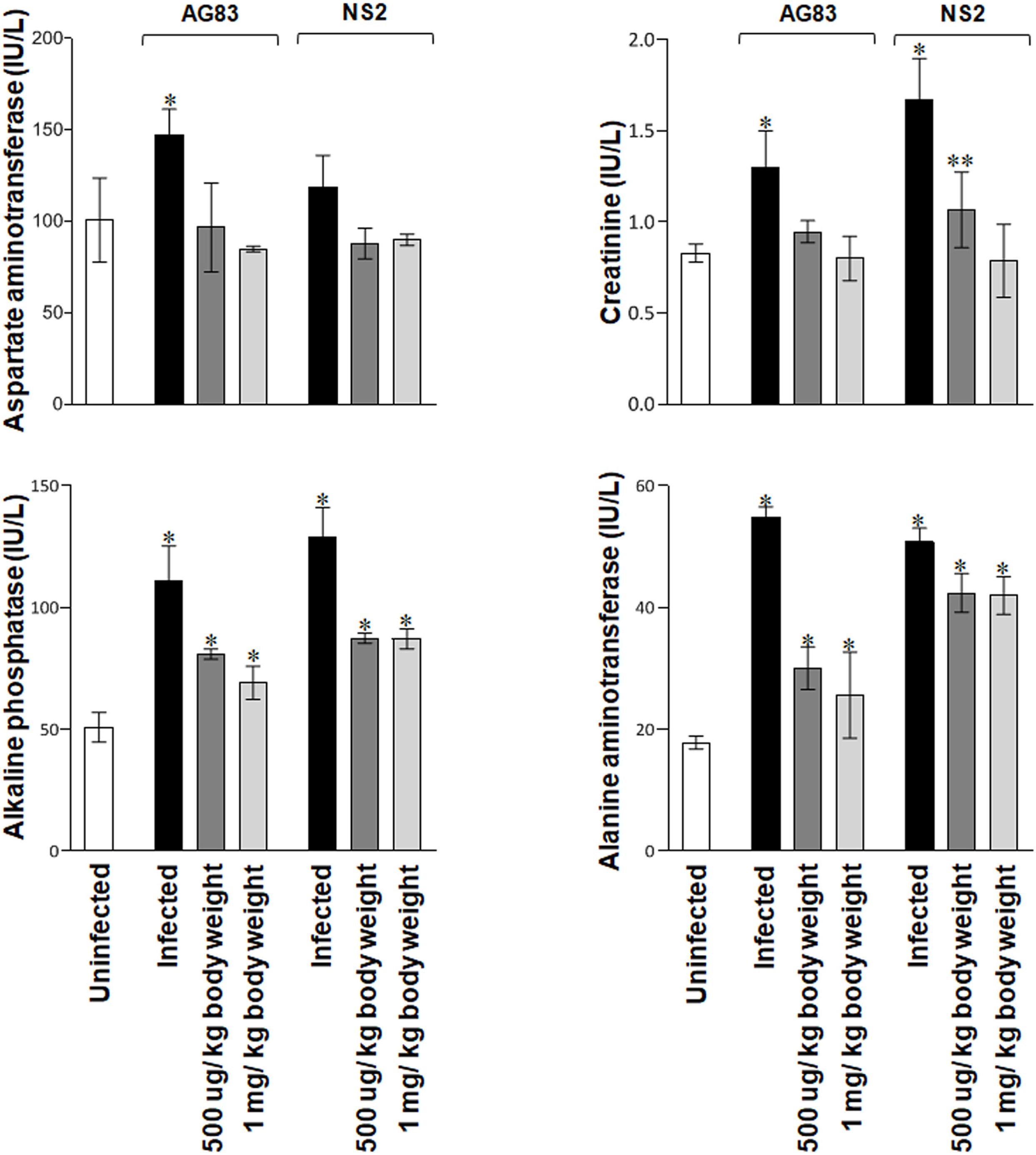
Estimation of the serum biomarkers (A) Aspartate transaminase, (B) Creatinine, (C) Alkaline phosphatase, and (D) Alanine transaminase levels from the *in vivo* experimental animals. All values are compared to the uninfected control group (*p<0.001, **p<0.004). Data represented as mean± SEM of two experiments, with 5 mice in each group.

### EL3 induced pro-inflammatory responses in the host, *in vivo*

Besides the disruption of the pro-parasitic thiol synthesis pathway, EL3 also induced the expression of the pro-inflammatory cytokines IL-12, TNF-α, IFN-γ, and IL-6 that instigate and amplify the innate immune responses in the host systems following parasite infection. TNF-α and IFN-γ were found up-regulated when the infected animals were treated with EL3, which are highly instrumental in generating nitric oxide in macrophages (21). EL3 also elevated the IL-6 dose-dependently, which is essential in resisting drug-resistant infection (Fig. 8) (22).

**Figure 8:**
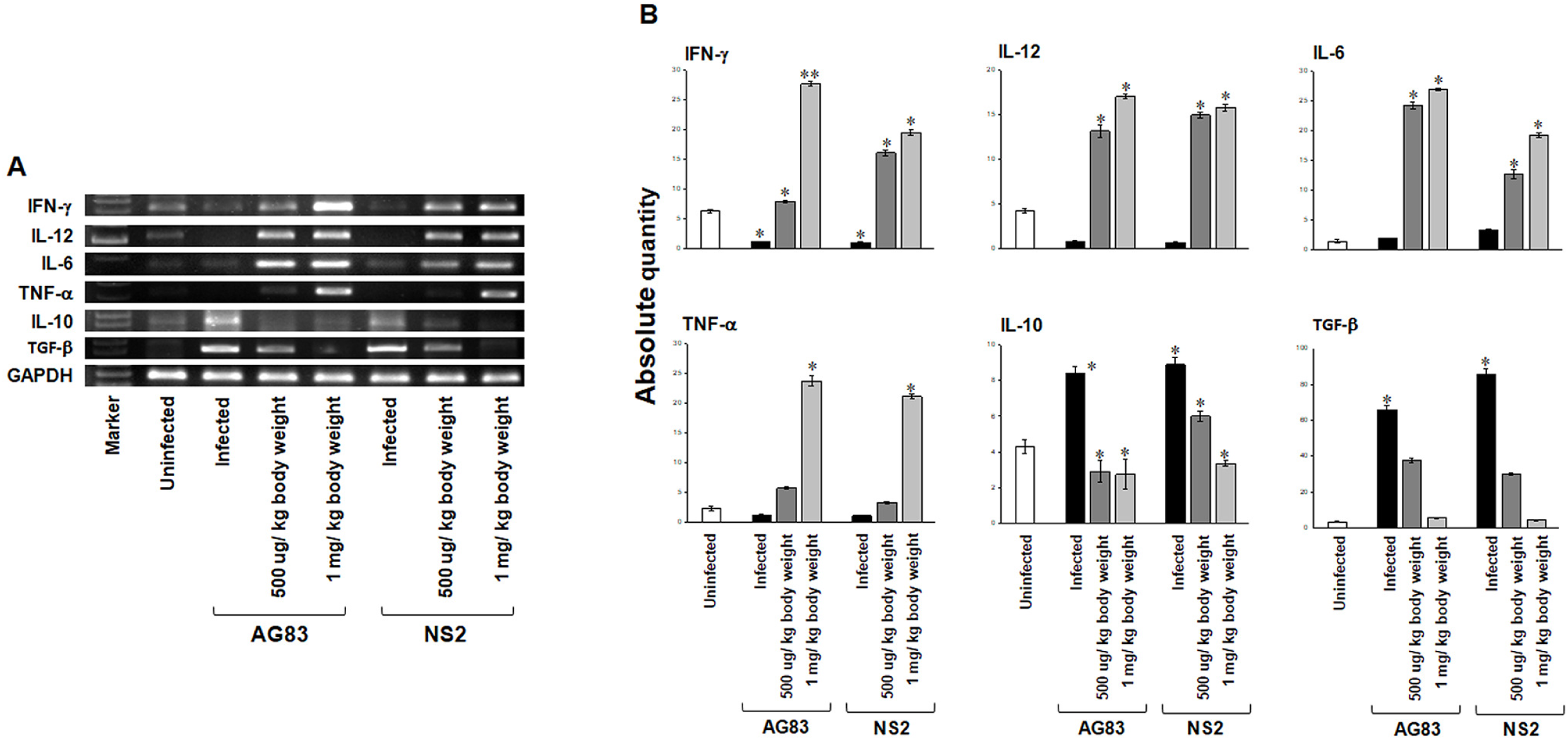
Induction of pro-inflammatory cytokines in the host, *in vivo*. (A) Induction of cytokines at the mRNA level. (B) Densitometry of the cumulative data, *p<0.001.All experiments were carried in duplicate, with 5 mice in each group.

### EL3 had a low serum bioavailability with maximum retention at 12 hours

The bioavailability of this compound in the serum of the host system was also checked at different time points, i.e., 1, 3, 6, 12, 24, 48, and 72 hours, post single intramuscular administration of 1 mg/kg body weight of the drug to naïve BALB/c mice. During the standardization process, the retention time for EL3 was initially set at 8.4 minutes at a wavelength of 275 nm. The resulting chromatogram showed no rise at the early 1-hour time point, but the concentration gradually increased; an initial concentration was detected at 3 hrs. post-injection, of 1.41 ± 0.1µg/ml (p<0.001 vs. 1 hr. mice serum) and 12 hours to a maximum of 6.8±0.2 µg/ml (p<0.001 vs. 1 hr. mice serum) post-intramuscular injection. The concentration then decreased to 1.3±0.1µg/ml (p<0.001 vs. untreated mice serum) at 48 hours for the untreated mice. No detection of EL3 was found at 1 hr and 72 hr. serum samples (Fig. 9, Table S13).

**Figure 9:**
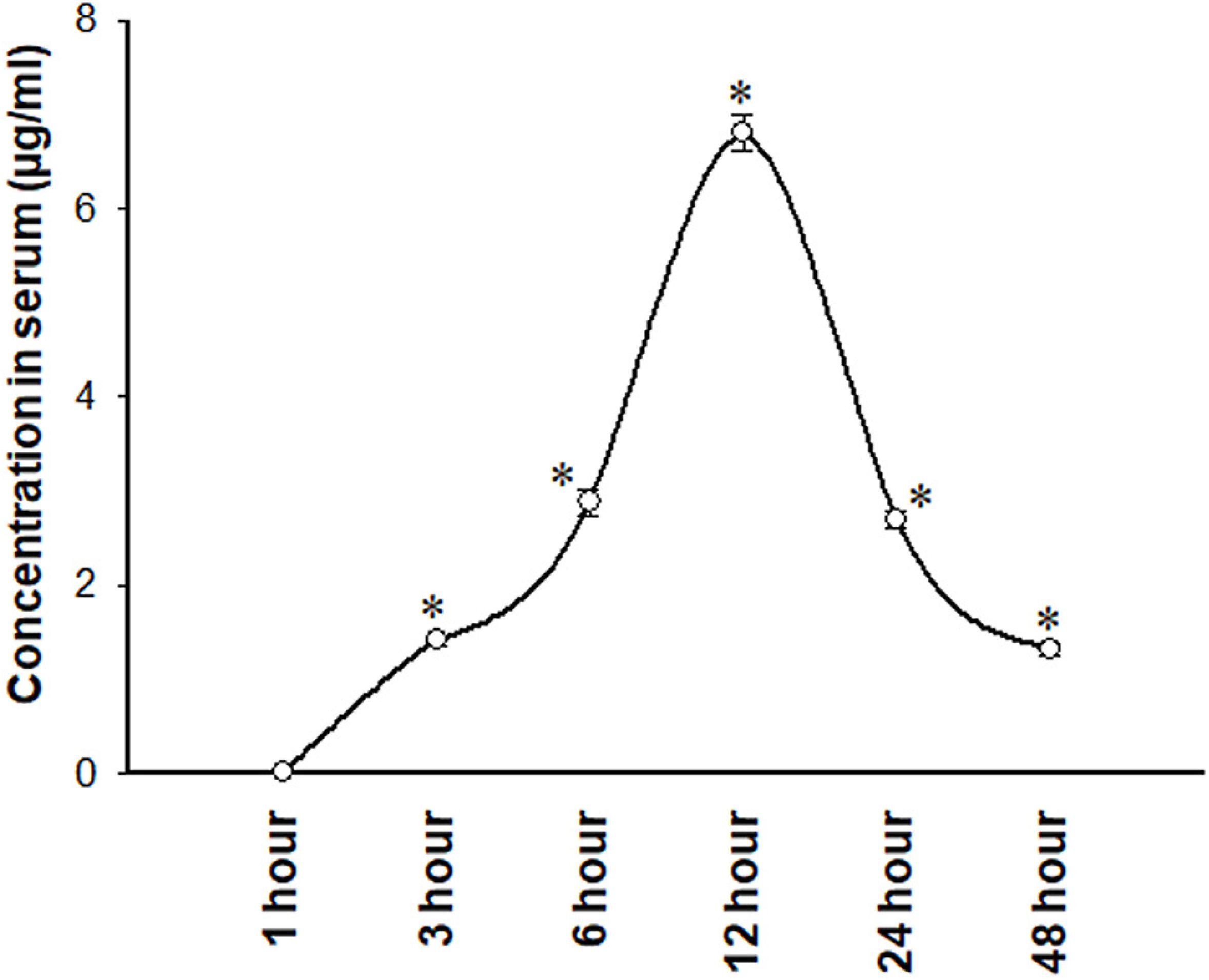
Bioavailability of EL3 from serum samples detected by HPLC at various time points (from 1h to 48h) post intramuscular injection of EL3 1 mg/kg b.w. dose *in vivo*. Significance was tested with respect to untreated mouse sera for various time point groups (*p<0.001). Data represented as mean± SEM of three different experiments.

## Discussion

The structure-activity relationship (SAR) investigation reveals that even minor modifications to the ethyl linoleate analogue (EL1) can significantly influence anti-leishmanial efficacy and pharmacokinetic behaviour. While increasing polarity by introducing additional hydroxyl groups in EL4, EL5, and EL6 enhanced aqueous solubility, this modification unexpectedly reduced bioactivity, likely due to impaired membrane permeability and diminished intracellular accumulation. In contrast, the mono-epoxygenated derivative EL3, which retains a balanced hydrophobic–hydrophilic character with fewer hydroxyl groups, exhibited the most promising *in vitro* and *in vivo* activity against both drug-sensitive and drug-resistant *L. donovani* strains. This enhanced efficacy is supported by its potent inhibition of promastigotes and amastigotes, low host toxicity, and its ability to downregulate key parasite survival enzymes involved in polyamine biosynthesis, particularly trypanothione reductase. Molecular docking and dynamic simulations further corroborate that EL3 forms stable hydrophobic interactions within the binding pockets of these enzymes, resulting in a higher binding affinity than its more polar counterparts. These findings, coupled with EL3’s favourable pharmacokinetic profile and its capacity to elicit a pro-inflammatory host response, underscore the importance of achieving an optimal balance between aqueous solubility and lipophilicity to maximize anti-leishmanial activity.

Drug design and therapeutic strategies against *L. donovani* infection neglect the difference in the pro-parasitic mechanisms that aid in escape from the host immune pathways, such as generating inflammatory cytokines or superoxides as ROS or NOS. This limits their use in various cases. Also, prevalent drugs in practice are reported to have severe toxicity and side effects on host metabolism. Longer persistence following a long-term treatment can lead to the emergence of drug-resistant strains that cause a higher rate of treatment failure. In this study, we isolated a bio-active ethyl linoleate from the ethyl acetate extract with anti-leishmanial activity and then further achieved a single epoxygenated derivative (EL3) from its synthetic analog (EL1), which had significant anti-proliferative properties against promastigotes and amastigotes of both the drug-resistant (NS2) as well as drug-sensitive (AG83) *L. donovani* parasites. Interestingly, the molecule was more effective against the resistant strain than the sensitive strain, both *in vitro* and *in vivo*. The inhibitory effect was due to the differential rates of gene expression of the thiol pathway between the virulent strains of the drug-resistant and drug-sensitive *L. donovani*. The enzymes γ-GCS, GS, and TR are needed in higher concentrations in resistant strains, and hence, lower expressivity of these enzymes marks the exceptional role of EL3 as an anti-leishmanial lead in therapeutics and drug design. Moreover, a secondary mode of action involving the induction of pro-inflammatory cytokine gene expression in macrophages also adds to the efficacy of the compound.

## Materials and methods

### Collection of Meripilus giganteus

The wild edible mushroom *M. giganteus* (Pers.) P. Karst, a Basidiomycota fungus, is found on stumps of freshly fallen trees and at the base of standing trees; often apparently growing from the ground, but always in contact with wood, widely distributed in the Northeastern and Eastern parts of the Himalayas (23). The basidiocarps were collected from different areas of Darjeeling during May to September, 2012-2015 (24). Information on edibility was gathered through discussions and direct interviews with local people and by direct observation of how this mushroom was collected and used. The damaged, infected, and very young fruit body of this mushroom was avoided, and the sample was collected preciously. The morphological and ecological features were noted, and color photographs of the materials were taken during field trips. After the specimens were brought to the laboratory, the macroscopic and microscopic properties were determined. Then the specimen was identified according to the previously described methods (23, 25, 26). The voucher specimen of the mushroom was deposited with the accession code CHU AM266 in the Mycological Herbarium of the Department of Botany, University of Calcutta (24).

### Isolation of active molecule and synthesis of derivatives

After the collection of the raw edible *M. giganteus* from the local market of Gangtok, India, the samples were dried in a hot air oven at 30°C for 48 hours. The dried mushroom was then ground into a fine powder using a mixer grinder to facilitate optimal metabolite extraction with different solvents. For bioassay-guided isolation of active compound(s), different crude extract was prepared using solvents of varying polarities to extract metabolites selectively. Initially, 20 g of finely powdered *M. giganteus* was soaked in 200 mL of petroleum ether in a round-bottom flask and gently stirred at room temperature for 48 hours to extract non-polar metabolites. The mixture was then filtered using vacuum filtration, and the solvent was evaporated at room temperature using a rotary evaporator, yielding a greasy brownish extract residue for bioassay. The remaining residue was subsequently soaked in 200 mL of chloroform under gentle stirring for 48 hours. Vacuum filtration followed by evaporation of solvent from the filtrate gives a yellowish, greasy material that has been used for bio-activity evaluation. The residue was subsequently treated with ethyl acetate (EtOAc) and methanol using a similar procedure, and the resulting solvent-extracted residues were prepared for bioactivity assays. These extracted residues were then evaluated for their bioactivity against drug-sensitive and drug-resistant promastigotes. Among them, the ethyl acetate extract of *M. giganteus* exhibited notably intriguing bioactivity against the drug-resistant strain of *L. donovani*. The significant *in vitro* bioactivity of the ethyl acetate-extracted residue against the drug-resistant strain of *Leishmania* prompted us to bioactivity-guided isolation of pure molecules from this extract.

The initial thin layer chromatographic (TLC) assessment of the residue of the ethyl acetate extract displayed the presence of multiple closely spaced spots under different staining agents (Fig. S1A). Due to the presence of various clusters of closely spaced spots in the TLC plate, we found it convenient to identify the bioactive cluster through initial preparative thin layer chromatographic fractionation of EtOAc extract. The fractionation was carried out using silica gel GF254 (Merk) on a 20 cm x 20 cm glass plate using Chloroform:Di-ethylEther: Pet Ether (10:2:1) as the optimized solvent system. Finally, we used an HPLC system equipped with reverse phase semi-preparative column (C18, Agilent, Column: Zorbax, 9 × 250 mm, particle size 5 μm, flow rate: 0.6 mL/min) and acetonitrile: water (7:3) as the optimized mobile phase to isolate the active compound from this bioactive fraction, obtaining the following chromatogram.

^1^H, ^13^C {1H}, NMR spectra were collected using Bruker Avance III 400 (^1^H: 400 MHz, ^13^C {1H}: 100 MHz) and were referenced to the resonances of the solvent used with TMS as internal standard. The chemical shifts were recorded in parts per million (ppm, δ) relative to CDCl_3_ (7.28 ppm for ^1^H and 77.00 for ^13^C) and DMSO *d_6_* (2.49 for ^1^H and 40.09 for ^13^C), and coupling constants (*J*) are reported in Hertz (Hz). Coupling patterns are indicated as: br (broad), s (singlet), d (doublet), t (triplet), q (quartet), p (pentet “quintet”), dd (doublet of doublet), td (triplet of doublets), or m (multiplet). Mass spectra were recorded on Accurate Mass Q-TOF LC/MS (Agilent Technologies Singapore, G6520B). Thin layer chromatography (TLC) was carried out to monitor the progress of the reactions using Merck pre-coated TLC plates (silica gel 60 F254 0.25 mm) and visualized by 254 nm and 366 nm UV light. Preparative thin-layer chromatography was carried out using silica gel GF254 (Merk) on a 20 cm x 20 cm glass plate. For isolation of active compound, we used HPLC system (Agilent, Column: Zorbax, 9 X 250 mm of particle size 5μm. Column chromatography was performed using silica gel (particle size 60-120 mesh) and eluted with a petroleum ether and ethyl acetate mixture. The structure and NMR spectra of the isolated peak 5 and synthetic derivatives have been provided (Fig. S2 - S7).

### Parasites and animals

Drug-sensitive AG83 parasites were originally obtained from CSIR-Indian Institute of Chemical Biology, Kolkata Jadavpur (10, 11), and drug-resistant NS2 parasites were kindly provided by Professor Mitali Chatterjee, Institute of PG Medical Education & Research, Kolkata, West Bengal, India (12, 13). *L. donovani* promastigotes were transformed from splenic aspirates of infected BALB/c mice as described previously (11). Male 4-5 weeks old BALB/c were procured from Centre for Laboratory Animal Research and Training, Kalyani, West Bengal, and allowed with rodent pellet diet, *ad libidum,* with a 12-hour cycle of light and dark as per CPSEA guidelines (File no. IAEC-1394/2015-16/5, dated 16.12.2015) following ARRIVED guidelines of the Institutional Animal Ethics Committee, WBSU, Barasat (13).

### Anti-promastigote activity and determination of inhibitory concentration, *in vitro*

The anti-promastigote activity of ethyl linoleate (Peak 5) and its synthetic derivatives (EL1 to EL6) was evaluated against the promastigotes of drug-sensitive and drug-resistant *L. donovani,* using the modified MTT assay, using the conventional tetrazoliumMTT salt (27). Briefly, 3×10^4^ promastigotes of both drug-sensitive and drug-resistant strains were cultured in each well of 96-well plates (purchased from Genaxy Scientific Pvt. Ltd), in 100 µl medium. After being treated with respective drugs for 48 hours, MTT was added at a 5 mg/ml concentration and allowed to form the formazan crystals, which were subsequently dissolved, and OD was taken using a multiplate reader (Bio-rad, USA).

### Anti-amastigote assay and evaluation of cytotoxicity in infected peritoneal macrophages, *in vitro*

Macrophages were obtained from 4% thioglycolate-stimulated peritoneal exudates of BALB/c mice, cultured for 48 hours for adhesion and extension. Further, the peritoneal macrophages were infected with *Leishmania* promastigotes (cells: parasite = 1:10) in 10% FBS supplemented RPMI 1640 (28). Infected macrophages were treated with 25, 50, and 75 µg/ml of isolated ethyl linoleate, synthetic ethyl linoleate (EL1), and its synthetic derivatives, EL2 and EL3, for 48 hours. Giemsa-stained micrographs were observed under a microscope (Carl Zeiss, Axioscope) to count the amastigotes/ 100 macrophages (6,7).

### *In vivo* efficacy and host toxicity of EL3

Male BALB/c mice (4-6 weeks, 5 mice per group) were infected with 2×10^7^ parasites of both AG83 and NS2 strains via intravenous (IV) route and treated with, 0.25 mg, 0.35 mg, 0.5 mg, and 1 mg/kg body weight of EL3 (one-month post-infection) via intramuscular (IM) route for an alternative 5 days, and the animals were sacrificed at one and half month of the post-infected period (scheme of the *in vivo* experiment illustrated in Fig. S8). Parasite survival and proliferation in the spleen and liver were calculated from the Giemsa-stained micrographs and expressed by Stauber’s formula (7).

### Nephro- and hepato-toxicity assessment

Experimental animals were anesthetized before sacrifice using diethyl ether. Blood was collected from the tail vein, and tubes were placed in a slanting position while the blood was allowed to form a clot for 45 min. Tubes were centrifuged at 1500Xg for 30 minutes, and the sera were collected from the supernatant without hemolysis and stored at -20°C for further use. Serum parameters for drug toxicity, such as creatinine, AST, AP, and ALT levels, were measured using Autospan kinetic assay kits, as per the manufacturer’s protocol (29).

### Bioavailability of EL3 in serum, *in vivo*

Blood was collected from naïve BALB/c mice (4-5 weeks) treated with a single intramuscular dose of 1 mg/kg body weight of EL3 and the serum was isolated at various time points (1, 3, 6, 12, 24, 48, and 72 hours post-treatment) were subjected to protein precipitation using acetonitrile (ACN) followed by centrifugation at 14000g for 15 minutes at 4°C. The clear supernatant from each sample was filtered using a 0.22 µm syringe filter before HPLC.

The presence of EL3 in the sera was detected in a C18 reverse phase column-Spherisorb (Waters) with dimensions of 4.6 x 150 mm and a 5 µm particle size. The separation was carried out using a mobile phase consisting of (a) HPLC-grade water and (b) HPLC-grade ACN in a gradient over 25 minutes at a flow rate of 1 ml/min. Dual absorbance detectors were used for detection, with one set at 275 nm and the other at 254 nm. Samples were injected using a Hamilton micro-syringe into the Shimadzu UFLC system (29).

### Analysis of intracellular *L. donovani* enzymatic pathway and host-specific pro-inflammatory cytokines, *in vivo*

4-6 weeks male BALB/c mice were divided into groups of uninfected, infected control (both AG83 and NS2), infected with AG83 and treated with 0.5 mg/kg body weight dose, infected with AG83 and treated with 1 mg/kg body weight dose, infected with NS2 and treated with 0.5 mg/kg body weight dose, infected with NS2 and treated with 1 mg/kg body weight dose of EL3. All doses administered intramuscularly, on each alternate day, up to 14 14-day dose regimen. The total mRNA was isolated from whole spleen tissue for real-time PCR of γ-GCS, GS, and TR transcripts from the intracellular parasites in a Bio-Rad thermo cycler using iTaq^TM^ SYBR green master mix. The analyses were performed using Bio-Rad CFX-maestro software. The relative mRNA expression (fold changes) was calculated using the 2^-(ΔΔCT)^ method (30).

Splenocytes from a similar set of animals were isolated and pulsed with 25 µg crude soluble antigen and maintained in RPMI 1640 medium with 5% CO_2_ concentration, at 37°C for 6 hours. The cDNAs were subsequently amplified with specific primers for IL-10, TGF-β, TNF-α, IFN-γ, IL-12, and IL-6 (Table S13) using a semi-quantitative reverse transcriptase PCR in a thermal cycler (Eppendorf, Germany). In all cases, the specific expressions of the cytokines were normalized against murine GAPDH using the housekeeping gene (22).

Three-dimensional structure prediction of γ-glutamyl cysteine synthetase, glutathione synthetase, and trypanothione reductase from *Leishmania donovani*.

The sequences of the three target proteins, γ-GCS, GS, and TR from *L. donovani* were retrieved from the UniProt database using the specific IDs Q67BG3, E9BBX9, and P39050, respectively. The three-dimensional structures of these proteins were generated using MODELLER 9v10 and AlphaFold2 Colab (31, 32). In MODELLER, a total of 100 models were generated, and the best one was chosen based on the DOPE score (33). AlphaFold2, on the other hand, computes the pLDDT and pTM scores to assess the accuracy of its predictions, with the top-ranked prediction by pLDDT used for further analysis (34, 35). The stereochemical qualities of the three protein models were validated by Ramachandran plot using PROCHECK (36). The two-dimensional (2D) chemical structure of the synthesized ligand was sketched using ChemDraw and was converted into the corresponding standard three-dimensional (3D) structure by using open Babel (37, 38).

### Molecular docking of EL3 with the three target proteins

To assess the binding efficiency of the EL3 molecule in predicting its interaction with the three target proteins of *L. donovani*, molecular docking was employed using LeDock software (39). All proteins and EL3 molecules were energy minimized before docking. The three target proteins were arranged in a cubic box with a grid point spacing of 0.3750 Å. The 100 docking conformations were generated for each protein-ligand complex. The binding conformations were assessed based on their binding energy (kcal/mol), and the conformation with the lowest binding energy (indicating the strongest binding affinity) for each protein-EL3 complex was selected for further analysis. The molecular interactions of protein-ligand complexes were generated by using PLIP (Protein-Ligand Interaction Profiler) (40).

### Molecular dynamics simulation of protein-EL3 complexes

In this study, molecular dynamics (MD) simulations of three *L. donovani* protein-EL3 complexes, γ-GCS, GS, and TR were performed to investigate their stability and molecular interaction mechanisms. GROMACS v 2023 and AMBER 99SB force field were used for the MD simulations study (41, 42). EL3 topology was generated using ACPYPE (AnteChamberPYthon Parser interface) for GROMACS compatibility (43). Each protein-EL3 system was solvated in a cubic water box with a TIP3P water model by maintaining periodic boundary conditions (PBC) (44, 45). All three protein-ligand complexes were neutralized with sodium and chloride ions. Energy minimization of each system was performed using the steepest descent until the maximum force was smaller than 1000 kJ/mol/nm (46). Each protein-EL3 complex was equilibrated with a 200ps isothermal-isochoric ensemble, NVT followed by a 200ps isothermal-isobaric ensemble NPT. These two equilibration methods stabilized systems at 310K and 1 bar pressure. Temperature and pressure coupling were controlled using the Berendsen thermostat and Parrinello-Rahman methods, respectively (47, 48). The Particle Mesh Ewald (PME) method was used to calculate the long-range electrostatic interactions, with cut-off radii of 0.9 nm set for both Van der Waals and short-range Coulombic interactions (49). The Linear Constraint Solver (LINCS) algorithm was used to fix the peptide bond lengths and angles, ensuring simulation stability and accurate dynamics of the protein-EL3 complexes (50). Production simulations were conducted for 100ns, with outputs saved at 10ps intervals. Principal Component Analysis (PCA) or essential dynamics was conducted on backbone atoms of three target proteins to identify primary motion patterns, with covariance matrices generated to examine system flexibility (51). For binding affinity assessments, the MM-PBSA (Molecular Mechanics Poisson-Boltzmann Surface Area) method was employed on selected trajectory frames of each system, calculating interactions like electrostatics and van der Waals forces, yielding detailed free energy profiles across the three protein-EL3 complexes (52). Stability metrics such as RMSD, RMSF, Rg, and SASA were analyzed to validate the reliability of the simulation outcomes.

### Estimation of binding free energy

To calculate the binding free energy of EL3 with the three target proteins, γ-GCS, GS, and TR from *L. donovani*, we employed the MM-PBSA method (52). Using the GROMACS tool, binding free energy calculations were estimated over the last 50 ns trajectory frames, ensuring the system had achieved equilibrium. A total of 100 snapshots were extracted from the trajectory at 0.5 ns intervals, providing representative conformations of the protein-EL3 complexes for reliable energy estimation.

### Statistical analysis

All results shown are representative of three different biological replicates of each experiment. The obtained data were analysed using Sigma Plot 11 software. Statistical analyses were done using ANOVA, and significance testing of each result was done by Tukey’s test. Results were represented as mean± SEM.

## Acknowledgments

We thank the Vice Chancellor of West Bengal State University for providing the research infrastructure required for this work. We acknowledge the DST-FIST, Govt. of India (Ref. SR/FST/LS1-001/2014), and DBT-BOOST, Govt. of West Bengal (Ref. 49[11]/BT [Estt]/1P-4/2013 [Part-1]), for providing funds for the real-time PCR facility in the Department of Zoology, WBSU, Barasat. We also acknowledge the iSTEM facility of the Bose Institute, Kolkata, for providing us with the HPLC facility. SC acknowledges CSIR-IICB for infrastructure support. AC acknowledges ICMR for research associateship [BMI/11(55)2022].

## APPENDIX

TLC, thin layer chromatography, HPLC-High performance liquid chromatography, DMSO-Dimethyl sulphoxide, MTT-3-(4,5-dimethylthiazol-2-yl)-2,5-diphenyl-2H-tetrazolium bromide, IL-Interleukin, IFN-Interferon,

## CRediT authorship contribution statement

**Supriya Nath:** Investigation, Methodology, Formal analysis, Validation, Writing – original draft; **Karan Chhetri:** Chemical synthesis, Methodology, Formal analysis; **Aabid Hussain:** Investigation, Methodology, Formal analysis, **Ankur Chaudhuri:** Investigation, Methodology, *In silico* Validation, Formal analysis; **Joydip Ghosh:** Investigation; **Sondipon Chakraborty:** Investigation, Methodology, Formal analysis, Validation; **Debarati Mukherjee:** Investigation; **Mintu Karan:** Investigation; **Bikramjit Raychaudhury:** Methodology; **Krishnendu Acharya:** Methodology; **Saikat Chakrabarti:** Writing**-**review & editing, Data curation, Conceptualization, Bioinformatics; **Biswajit Gopal Roy:** Writing-review & editing, Data curation, Conceptualization, Chemical synthesis, Supervision**; Chiranjib Pal:** Writing-review & editing (Final draft), Validation, Project administration, Fund acquisition, Data curation, Conceptualization, Supervision.

